# Free fatty acid receptor 4 in cardiac myocytes ameliorates ischemic cardiomyopathy

**DOI:** 10.1101/2024.04.12.589280

**Authors:** Michael J. Zhang, Sergey Karachenets, Dylan J. Gyberg, Sara Puccini, Chastity L. Healy, Steven C. Wu, Gregory C. Shearer, Timothy D. O’Connell

**Author notes:** Correspondence to: Timothy D. O’Connell, PhD, Department of Integrative Biology and Physiology University of Minnesota School of Medicine, 3-141 CCRB, 2231 6th Street SE Minneapolis, MN 55414 -and-Gregory C. Shearer, PhD Department of Nutritional Sciences 110 Chandlee Laboratory University Park, PA 16802.

## Abstract

**Aims:** Free fatty acid receptor 4 (Ffar4) is a receptor for long-chain fatty acids that attenuates heart failure driven by increased afterload. Recent findings suggest that Ffar4 prevents ischemic injury in brain, liver, and kidney, and therefore, we hypothesized that Ffar4 would also attenuate cardiac ischemic injury.

**Methods and Results:** Using a mouse model of ischemia-reperfusion (I/R), we found that mice with systemic deletion of Ffar4 (Ffar4KO) demonstrated impaired recovery of left ventricular systolic function post-I/R with no effect on initial infarct size. To identify potential mechanistic explanations for the cardioprotective effects of Ffar4, we performed bulk RNAseq to compare the transcriptomes from wild-type (WT) and Ffar4KO infarcted myocardium 3-days post-I/R. In the Ffar4KO infarcted myocardium, gene ontology (GO) analyses revealed augmentation of glycosaminoglycan synthesis, neutrophil activation, cadherin binding, extracellular matrix, rho signaling, and oxylipin synthesis, but impaired glycolytic and fatty acid metabolism, cardiac repolarization, and phosphodiesterase activity. Kyoto Encyclopedia of Genes and Genomes (KEGG) pathway analysis indicated impaired AMPK signaling and augmented cellular senescence in the Ffar4KO infarcted myocardium. Interestingly, phosphodiesterase 6c (PDE6c), which degrades cGMP, was the most upregulated gene in the Ffar4KO heart. Further, the soluble guanylyl cyclase stimulator, vericiguat, failed to increase cGMP in Ffar4KO cardiac myocytes, suggesting increased phosphodiesterase activity. Finally, cardiac myocyte-specific overexpression of Ffar4 prevented systolic dysfunction post-I/R, defining a cardioprotective role of Ffa4 in cardiac myocytes.

**Conclusions:** Our results demonstrate that Ffar4 in cardiac myocytes attenuates systolic dysfunction post-I/R, potentially by attenuating oxidative stress, preserving mitochondrial function, and modulation of cGMP signaling.

## Introduction

Coronary heart disease (CHD) is the leading cause of death globally, resulting in an estimated 9.4 million deaths worldwide in 2021.^1^ Although effective and timely coronary revascularization has led to a decrease in the incidence rates of CHD mortality in resource rich countries, the incidence rates of heart failure (HF) from ischemic cardiomyopathy (ICM) continue to rise.^2^ Despite the necessity of restoring blood flow, myocardial reperfusion can lead to severe injury. Principally, ischemia-reperfusion (I/R) impairs the mitochondrial respiratory chain, increases mitochondrial ROS generation, and induces irreversible mitochondrial damage and opening of the mitochondrial permeability transition pore, ultimately leading to necrotic and apoptotic cardiac myocyte cell death.^3^ Cardiac myocyte death leads to the release of damage associated molecular patterns (DAMPs), activation of pattern recognition receptors (PRR), local production of cytokines/chemokines and infiltration of inflammatory cells 1-3 days following injury, initially neutrophils in the early inflammatory phase, followed by macrophages in the inflammatory/proliferative phases 3-7 days following injury.^4, 5^ Activation of PRRs also promotes NFκB signaling and upregulation of the NLRP3 inflammasome leading to further cardiac damage.^6^ However, therapeutically reducing oxidative stress and mitochondrial damage remains an unmet challenge.

Free fatty acid receptor 4 (Ffar4, GPR120) is a G-protein coupled receptor (GPCR) for long chain fatty acids, including cardioprotective μ3-polyunsaturated fatty acids (μ3-PUFAs), that attenuates metabolic dysfunction and resolves inflammation.^7, 8^ Ffar4 is expressed in many tissues including intestinal I, K, and L enteroendocrine cells, pancreatic α, β, and δ-cells, brain, and lung. More importantly, Ffar4 is also expressed in both cardiac myocytes and fibroblasts,^9, 10^ as well as macrophages,^11, 12^ indicating a potential role in the response to ischemic injury. Interestingly, accumulating evidence suggests that Ffar4 can mitigate ischemia/reperfusion (I/R) injury. In the brain, Ffar4 expression in microglia is increased after ischemic injury, and knockdown of Ffar4 abolished the neuroprotective effect of μ3-PUFAs.^13^ In Kupffer cells, activation of Ffar4 by μ3-PUFAs or the Ffar4 agonist GW9508 prevented ischemia-induced necrosis and elevation of liver enzymes.^14^ In the kidney, the Ffar4 agonist TUG-891 alleviated I/R-induced kidney injury, in part by activating an AMPK/SirT3 signaling pathway, thus inhibiting tubular epithelial cell senescence.^15^ Interestingly, Ffar4 also attenuates ROS production in cultured macrophages,^16, 17^ and in cultured hepatocytes, Ffar4 promotes mitophagy to protect against oxidative stress.^18^ Finally, Ffar4 prevents NLRP3 inflammasome activation in Kupffer cells ^19^ and macrophages.^20^ Collectively, these studies suggest that Ffar4 might attenuate oxidative stress to attenuate ischemic injury.

Previously, we demonstrated that Ffar4 attenuated pathologic remodeling secondary to pressure overload and cardiometabolic disease.^10, 21^ Recognizing the cardioprotective nature of Ffar4, combined with the ability of Ffar4 to attenuate oxidative stress and reduce ischemic injury in other organs, we hypothesized that in cardiac myocytes, Ffar4 would attenuate I/R injury. Briefly, our results indicate that in mice with systemic deletion of Ffar4 (Ffar4KO mice), loss of Ffar4 impaired the recovery of systolic function post-I/R in both male and female mice, without affecting initial infarct size. Transcriptome analysis three days post-I/R suggested deficits in mitochondrial metabolism and AMPK signaling, as well as augmentation of inflammatory processes and cellular senescence amongst other pathways in the infarcted myocardium from Ffar4KO mice that might have contributed to worsened outcomes. Loss of Ffar4 also increased phosphodiesterase 6c expression (PDE6c), a cyclic GMP-hydrolyzing phosphodiesterase previously uncharacterized in the heart whose expression is increased in human ischemic cardiomyopathy. Finally, we found that cardiac myocyte-specific overexpression of Ffar4 prevented systolic dysfunction post-I/R, indicating a direct cardioprotective effect of Ffar4 in cardiac myocytes. In summary, our data suggest that Ffar4 in cardiac myocytes attenuates I/R injury, preventing contractile dysfunction, possibly by attenuating oxidative stress, preserving mitochondrial function, and modulation of cGMP signaling.

## Methods

### Mice

Mice with systemic deletion of Ffar4 (Ffar4KO mice) were generated from cryopreserved sperm from C57Bl/6N-*Ffar4*^tm1^(KOMP)^Vlcg^ (Design ID: 15078; Project ID: VG15078) purchased from the Mutant Mouse Resource and Research Centers (MMRRC), UC Davis (Davis, CA, USA). Ffar4KO mice were backcrossed into the C57Bl/6J strain, and mice used in this study were from the 15-17th backcross generation.^10^

Mice with cardiac myocyte-specific overexpression of Ffar4 (CM-Ffar4-Tg) were developed using the CAG-CAT system (Plasmid: 53959, Addgene, Watertown, MA, USA).^22^ Briefly, mice have a stop sequence flanked by loxP sites (loxP-stop-loxP) upstream of the human Ffar4 cDNA sequence (NM 181745, OriGene Technologies Inc., Rockville, MD, USA), both under control of the cytomegalovirus (CMV) early enhancer element and chicken beta-actin (CAG) promotor. To activate Ffar4 expression, mice were infected with an AAV9-cTnt-CRE virus (10^12^ genome copies) or a control AAV9-cTnt-GFP virus via retro-orbital intravenous injection (Vector Biolabs, Malvern PA, USA).^23^ To validate Ffar4 overexpression, cardiac myocytes and non-myocytes were isolated 4 weeks after injection, and Ffar4 expression in both fractions was quantified by qRT-PCR as described below. Of note, we previously demonstrated that the commercially available antibodies to Ffar4 are non-specific,^24^ a consistent problem with antibodies to GPCRs, thus limiting our ability to detect Ffar4 protein.

### Transient coronary artery ligation (CAL) as a model of cardiac ischemia-reperfusion (I/R) injury

Mice were subjected to transient coronary artery ligation surgery to induce I/R injury as we previously described, which includes a full description of the method along with video tutorials.^25^ WT and Ffar4KO, male and female mice were subjected to CAL surgery at 12 weeks of age. CAG-CAT-Ffar4 mice, were injected with virus (AAV9-cTnt-GFP or AA9-cTnt-Cre) at 8 weeks of age, and underwent CAL surgery at 12 weeks of age. Briefly, mice were anesthetized with isoflurane (4% for induction, 2% for maintenance, 100% O_2_, 2 L/min), placed on a heated surface to maintain body temperature (monitored by rectal probe), and the chest was cleaned with betadine and 70% ethanol. A 1.5 centimeter vertical incision was made in the skin along the left midclavicular line. A horizontal mattress stitch with 6-0 silk suture was placed around this incision, without pulling it closed. A 1 cm section of skin superior to the skin incision was separated and an 8-0 nylon suture was tunneled underneath the skin. This leaves the loosening end of the coronary artery ligation suture exposed near the mouse’s neck, to prevent the mouse from pulling on the suture and to aid subsequent reperfusion. The pectoralis major and minor muscles were bluntly separated, and subsequently the pectoralis minor was separated from the ribcage. Using a hemostat, the thorax at the third intercostal space just medial to the left lung was bluntly pierced and the third and fourth rib were spread by approximately 5 mm. Upward pressure was applied on the thorax while pushing leftward on the right chest to exteriorize the heart. Using the pre-placed 8-0 suture, the heart was pierced at the base of the left ventricle immediately inferior to the left atrium (at the junction between the left ventricle and the pectoralis minor muscle), and returned the heart to the thorax, a slip knot was secured with each end of the 8-0 suture exteriorized, and the chest is closed with the original mattress suture. The pneumothorax was evacuated by placing gentle bilateral pressure on the mouse thorax, the isoflurane was discontinued, buprenorphine (1 mg/kg) was administered, and the mouse was removed from restraint and allowed to regain consciousness. After 60 min (WT vs Ffar4KO) or 120 min (WT vs CM-Ffar4-Tg) the mouse was anesthetized (3% for induction, 1.5% for maintenance, 100% O_2_, 2 L/min), the quick release knot is released, and the mouse is allowed to regain consciousness. Sham surgeries were performed, which consisted of all procedures except the ligation step.

### Echocardiography

Cardiac function was measured using a Vevo 2100 Imaging System (VisualSonics, Toronto, Ontario, Canada) equipped with an 18-38 mHz linear array transducer. Mice were anesthetized with isoflurane that was adjusted to maintain a heart rate between 400-500 beats per minute. Time-gated 2D cine loops were acquired from parasternal long axis, parasternal short axis and apical four chamber views. Left ventricular (LV) systolic function and LV volumes were quantified averaging ejection fraction and ventricular volumes obtained from 2D edge-tracking (LV Trace) and Simpson’s method of disks.^26^

### Measurement of ischemic area (IA) and area-at-risk (AAR)

IA and AAR were measured 24 h post-I/R.^25, 27^ Mice were anesthetized with isoflurane (3%), the original ligation was re-tied, and the heart perfused with 1% Evans Blue Dye (EBD) to demarcate the risk region by staining uninjured nonischemic tissue. The heart was then sliced into 2 mm sections and incubated in 1% triphenyltetrazolium chloride (TTC) at 37°C for 15 min to stain AAR (the infarct remains unstained). AAR is measured by EBD negative area, and infarct size is measured by TTC negative area.

### RNAseq

Three days after CAL or Sham surgery (I/R: WT n=8, Ffar4KO n=7; Sham: WT n=5, Ffar4KO n=5), hearts were removed, perfused free of blood, and in hearts from mice that underwent I/R surgery, the infarct region was separated from the non-infarct region, whereas sham hearts were left intact. RNA was isolated from all tissues using RNeasy Fibrous Tissue Mini Kit (Qiagen). 125bp paired end sequencing was performed using the HiSeq 2500 sequencer (Illumina) by the University of Minnesota Genomics Center. Data were analyzed by the University of Minnesota Informatics Institute. 2 × 125bp FastQ paired end reads (n=8.4 million average per sample) were trimmed using Trimmomatic (v 0.33) enabled with the optional “-q” option; 3bp sliding-window trimming from 3’ end requiring minimum Q30. Quality control on raw sequence data for each sample was performed with FastQC. Read mapping was performed via Hisat2 (v2.1.0) using the mouse genome (mm10) as reference. Gene quantification was done via Cuffquant for FPKM values and Feature Counts for raw read counts. Principal component analyses used CPMs that were filtered based on gene size (excluding genes less than 200bp) and variance less than 1 in raw read counts and the top 2,000 differentially expressed genes were included in the analysis (Prism 10.0). Data was subsequently analyzed using the BioJupies analysis tool,^28^ and Excel files for the analysis of differentially expressed genes, gene ontology, pathway analysis, transcription factor enrichment analysis, and kinase enrichment analysis are available. Kyoto Encyclopedia of Genes and Genomes (KEGG) pathway analysis of the top 3,000 differentially expressed genes comparing the infarcted myocardium from WT and Ffar4KO hearts was performed using Pathview.^29^

### Isolation and culture of adult cardiac myocytes

We previously published a comprehensive step-by-step procedure for the isolation and culture of adult mouse cardiac myocytes that was used in this study.^30^

### qRT-PCR

RNA was isolated from hearts using the RNeasy Fibrous Tissue Mini Kit (Qiagen, Germantown MD, USA). RNA concentration was determined by NanoDrop Spectrophotometer (Thermo Fisher, Waltham, MA, USA) and cDNA was synthesized by reverse transcription using the Bio-Rad iScript cDNA Synthesis Kit (Hercules, CA, USA). Gene expression was quantified by qRT-PCR using the Bio-Rad iTaq Universal SYBR Green SuperMix and Bio-Rad CFX96 Real-Time System.

Primers: (5’ to 3’)

Ffar4 For: CGG CGG GGA CCA GGA AAT, Rev: GTC TTG TTG GGA CAC TCG GA PDE6c For: GCG GCA GTT TGA AAC GGT G Rev: CAT CAT AGG CTG ACT CTG CAC

### Western blotting

Cardiac PDE6c protein expression was quantified by Western blot using a polyclonal antibody generated from rabbits immunized with a synthetic peptide between 271-300 amino acids from the central region of human PDE6c (SAB1301616, MilliporeSigma, St. Louis, MO, USA). Left ventricular tissue from WT or Ffar4KO mice was harvested, snap frozen and homogenized in RIPA Lysis Buffer (Source) using hard tissue homogenization beads (Benchmark Industrial, Sayreville, New Jersey, USA). After normalization of protein concentration by a Bradford protein assay, protein lysates were denatured in Laemmli buffer and separated by SDS-PAGE. Proteins were then transferred onto a polyvinylidene fluoride membrane. The membrane was blocked with 5% non-fat dry milk in phosphate buffered saline and incubated serially with primary and HRP-conjugated secondary antibodies. Blots were developed using an electrochemiluminescence kit (Bio-Rad, Hercules, CA, USA) and imaged using a Bio-Rad Chemidoc CCD camera. Densitometry was then performed using ImageJ.

### cGMP ELISA

Cardiac myocyte cGMP levels were quantified using the Cyclic GMP ELISA Kit (Cayman Chemical, Ann Arbor, MI, USA) according to the manufacturer’s instructions. All samples were acetylated to increase assay sensitivity. Atrial natriuretic peptide (Cat# 24276), vericiguat (Cat# 35253), and TUG-891 (Cat# 17035) were also purchased from Cayman Chemical. For quantification of cardiac myocyte PDE6c activity, isolated adult mouse cardiac myocytes were stimulated with vehicle, 10 µM atrial natriuretic peptide, or 10 µM vericiguat for 30 minutes before cell lysis and harvest for cGMP ELISA. For quantification of cGMP levels after TUG-891 stimulation *in vivo*, mice were injected IP with 35 mg/kg of TUG-891 (100 µL injection volume, corn oil vehicle) every 24 hours for 3 days. Cardiac myocytes were then isolated for cGMP quantification by ELISA.

### Statistics

All data are presented as mean ± SEM, and a Shapiro-Wilk test was used to test for a normal distribution. Comparisons of two groups were performed using a Student’s t-test, whereas comparisons of multiple groups were performed using a two-way ANOVA with repeated measures and a Tukey’s post-test with Prism 10.0 (GraphPad Prism, San Diego, CA). P<0.05 was considered significant. Results of post-test comparisons are only shown when the primary interaction was significant.

## Study Approvals

### Animal

All procedures on animals conformed to the NIH Guide for the Care and Use of Laboratory Animals and were reviewed and approved by the Institutional Animal Care and Use Committee at the University of Minnesota.

### Additional compliance statements

For all surgeries, mice were anesthetized by induction with 3% isoflurane, and once anesthesia was induced, mice were maintained at 1.5% isoflurane, and verified by toe-pinch. Post-surgery, buprenorphine-sustained release (1 mg/kg SC) was administered for pain management during the first 24 hours post-surgery and as needed thereafter. At the indicated time points post-surgery or for the isolation of cardiac myocytes, mice were anesthetized with 3% isoflurane, verified by toe-pinch, followed by removal of the heart in accordance with recommendations from the American Veterinary Medical Association. Finally, the data underlying this article are available in the article and in its online supplementary material.

## Results

### Loss of Ffar4 worsens left ventricular systolic dysfunction induced by ischemia-reperfusion (I/R) injury in male and female mice

To test the hypothesis that Ffar4 is required for an adaptive response to cardiac injury induced by I/R, we quantified left ventricular remodeling in male wild-type (WT) mice and mice with systemic deletion of Ffar4 (Ffar4KO) subjected to 60 minutes of ischemia by transient coronary artery ligation (CAL) followed by reperfusion for four weeks. After three days, I/R induced a similar and significant decrease in left ventricular systolic function measured by ejection fraction (EF) in both male WT and Ffar4KO mice (Figures 1A-C). From that point forward, systolic function (EF) recovered to almost baseline values in male WT mice by four weeks, but remained suppressed in male Ffar4KO mice (Figure 1A). Additionally, neither left ventricular weight (Figure 1D) nor fibrosis (Figures 1E-F) were different in WT versus the Ffar4KO hearts after four weeks. In a separate cohort of mice, we assessed the extent of damage induced by I/R after 24 hours. Neither area-at-risk nor infarct area were different in male WT versus the Ffar4KO hearts (Figures 1G-I).

**Figure 1.**
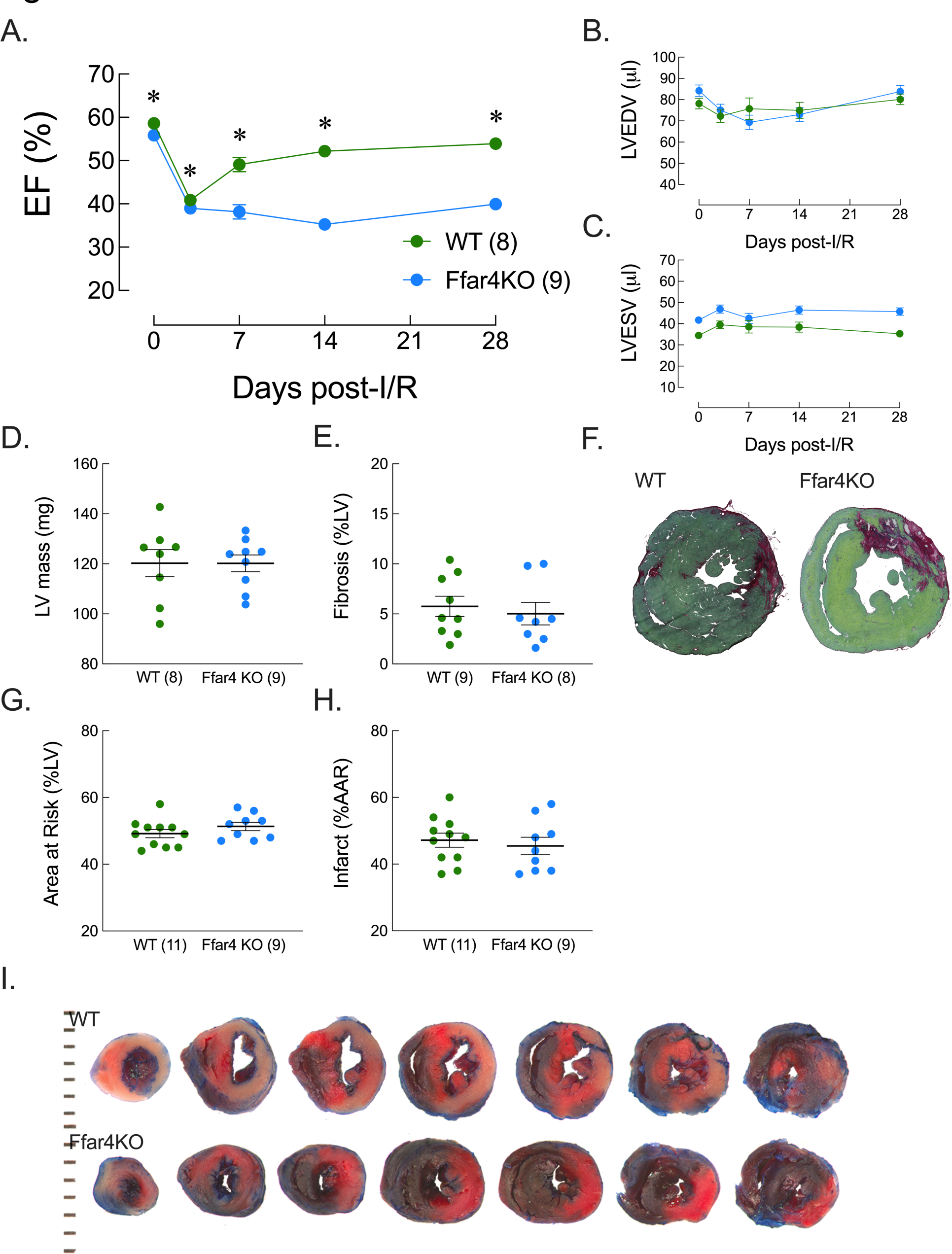
Loss of Ffar4 worsens left ventricular systolic dysfunction induced by ischemia-reperfusion (I/R) injury in male mice. Male WT and Ffar4KO mice were subjected to 60 min of ischemia followed by reperfusion for 4 weeks. Systolic function and ventricular volumes were measured by echocardiography from 2D-parasternal long-axis views prior to I/R surgery, and 3, 7, 14, and 28 days post-ischemia. (**A**) Ejection fraction, EF; (**B**) Left ventricular end diastolic volume, LVEDV; (**C**) Left ventricular end systolic volume, LVESV; and (**D**) Left ventricular mass measured by echocardiography at 4 weeks. After 4 weeks, ventricular fibrosis was measured by Sirius red-fast green staining (SR/FG). (**E**) Ventricular fibrosis; and (**F**) Representative ventricular sections stained with SR/FG. In a separate cohort, area-at-risk and infarct size were measured 24 hr post-I/R using triphenyltetrazolium chloride (TTC) staining. (**G**) Area-at-risk; (**H**) Infarct area; and (**I**) Representative TTC staining of WT and Ffar4KO hearts, serial section starting with the apex on the left to the base on the right. In A-E and G and H, data are expressed as mean±SEM. In A-C, data were analyzed by two-way ANOVA with repeated measures with a Tukey post-test to detect differences between groups. In D, E, G, and H groups were compared by student’s t-test. P<0.05 was considered significant.

Similar results were observed in female mice. After three days, I/R induced significant left ventricular systolic dysfunction (EF) in female WT and Ffar4KO mice, and while systolic function recovered in female WT mice by four weeks, it remained depressed in female Ffar4KO mice (Figures 2A-C). Further, neither left ventricular weight nor fibrosis were different after four weeks in female WT and Ffar4KO mice (Figures 2D-E). Again, in a separate cohort of mice neither area-at-risk and infarct area were different after 24 hours (Figures 2F-G). These results indicate that loss of Ffar4 impairs recovery of systolic function post-ischemia in both male and female mice.

**Figure 2.**
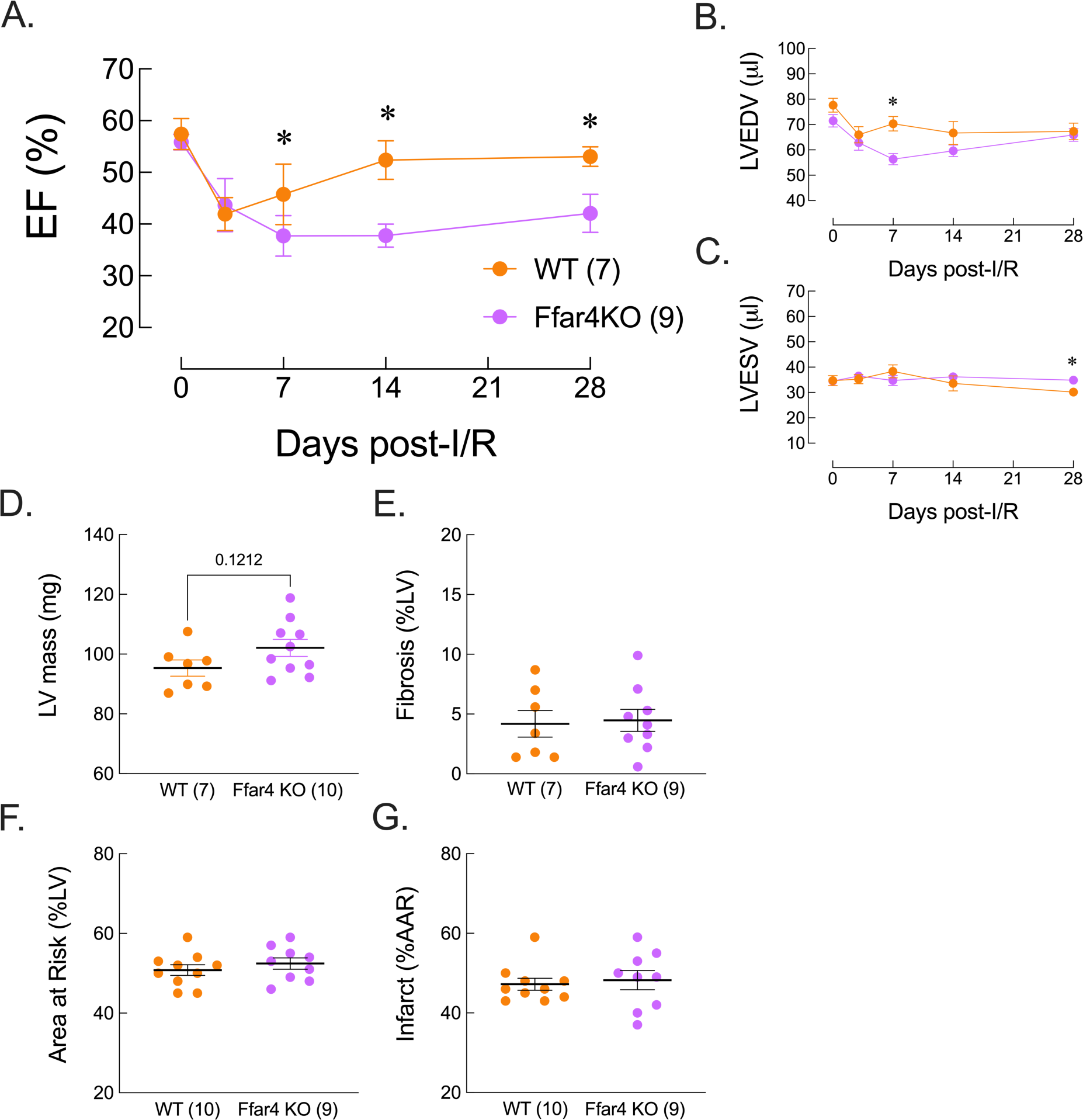
Loss of Ffar4 worsens left ventricular systolic dysfunction induced by Ischemia-reperfusion (I/R) injury in female mice. Similar to studies in males, female WT and Ffar4KO mice were subjected to 60 min of ischemia followed by reperfusion for 4 weeks. Systolic function was measured as in Figure 1. ((**A**) Ejection fraction, EF; (**B**) Left ventricular end diastolic volume, LVEDV; (**C**) Left ventricular end systolic volume, LVESV; and (**D**) Left ventricular mass measured by echocardiography at 4 weeks. (**E)** After 4 weeks, ventricular fibrosis was measured by Sirius red-fast green staining (SR/FG). In a separate cohort, area-at-risk and infarct size were measured 24 hr post-I/R using triphenyltetrazolium chloride (TTC) staining. (**F**) Area-at-risk; (**G**) Infarct area. In A-G, data are expressed as mean±SEM. In A-C, data were analyzed by two-way ANOVA with repeated measures with a Tukey post-test to detect differences between groups. In D-G, groups were compared by student’s t-test. P<0.05 was considered significant.

### Transcriptome analysis of the infarcted myocardium

Contractile function is unaffected in Ffar4KO mice at baseline, and there is no evidence that Ffar4 regulates chronotropic or inotropic responses in the heart. Therefore, to uncover potential mechanistic explanations for the worsened systolic function observed in Ffar4KO mice post-I/R, we performed a transcriptome analysis on infarcted heart by bulk-RNAseq three days post-I/R. Specifically, we compared the infarcted myocardium in male WT and Ffar4KO hearts, using the non-infarcted myocardium as controls (Figure 3A). A principle component analysis (PCA) of the top 2,000 differentially expressed genes identified global patterns of differential gene expression (Figure 3B). There was considerable overlap in the transcriptomes from the non-infarcted myocardium in WT and Ffar4KO hearts. Importantly, the transcriptome in the infarcted myocardium from WT hearts showed significant variability along PC2 relative to the non-infarcted myocardium from both WT and Ffar4KO hearts, which was further magnified in the infarcted myocardium from Ffar4KO hearts.

**Figure 3.**
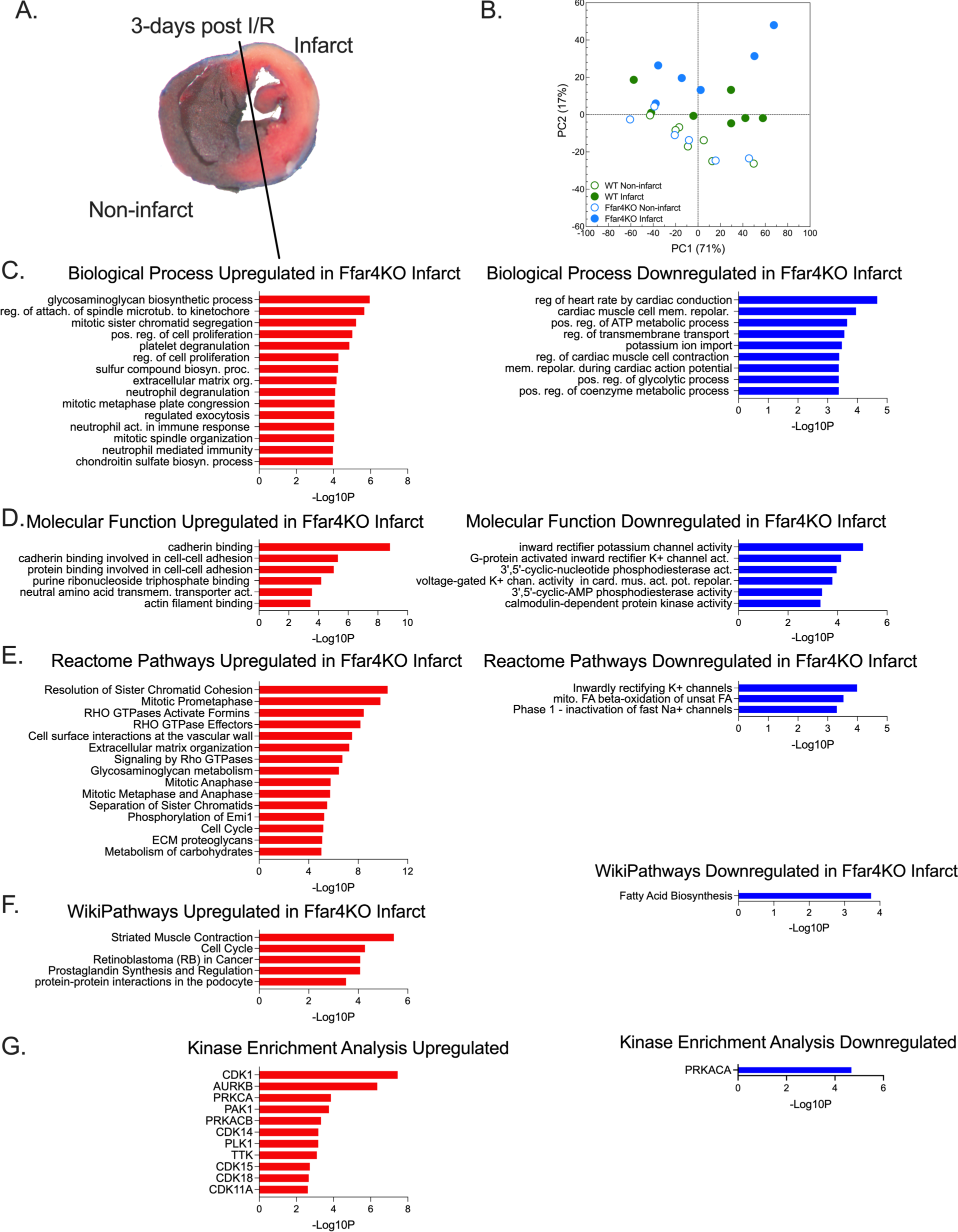
Transcriptome analysis of the infarcted myocardium. (**A**) Male WT and Ffar4KO mice were subjected to 60 min of ischemia followed by reperfusion for 3 days, at which point the ventricle was divided into infarcted and non-infarcted regions and total RNA was isolated and submitted for bulk-RNAseq. (**B**). A principle component analysis (PCA) was performed on the top 2,000 differentially expressed genes with PC1 (71% of the variability) plotted on the x-axis and PC2 (17% of the variability) plotted on the y-axis. Using BioJupies analysis software,^28^ changes in gene ontology (GO) (**C**) Biological process and (**D**) Molecular function, as wells as (**E**) Reactome Pathways, (**F**) Wikipathways, and (**F**) Kinase Enrichment were analyzed. The top 15 categories significantly changed categories or all categories with P<0.0005 (if less than 15) were plotted as −log10P values on the x-axis and GO categories on the y-axis.

To compare the transcriptomes of the infarcted myocardium from WT and Ffar4KO hearts, we performed a gene ontology (GO) and pathway enrichment analysis using BioJupies analysis software.^28^ (Figure 3C-G). This analysis identified significantly enriched (red) or diminished (blue) gene ontologies (Figure 3C, Biological Process; Figure 3D, Molecular Function), or pathways (Figure 3E, Reactome Pathways; Figure 3F, WikiPathways; Figure 3G, Kinase Enrichment Analysis). In aggregate, glycosaminoglycan biosynthesis, particularly chondroitin biosynthesis, neutrophil activation, cadherin binding, extracellular matrix, rho signaling, and oxylipin (prostaglandin) synthesis pathways were enriched in the Ffar4KO infarcted myocardium (Table 1). Conversely, cardiac myocyte repolarization, glycolytic metabolism, fatty acid metabolism, and phosphodiesterase pathways were diminished in the Ffar4KO infarcted myocardium (Table 2).

**Table 1:**
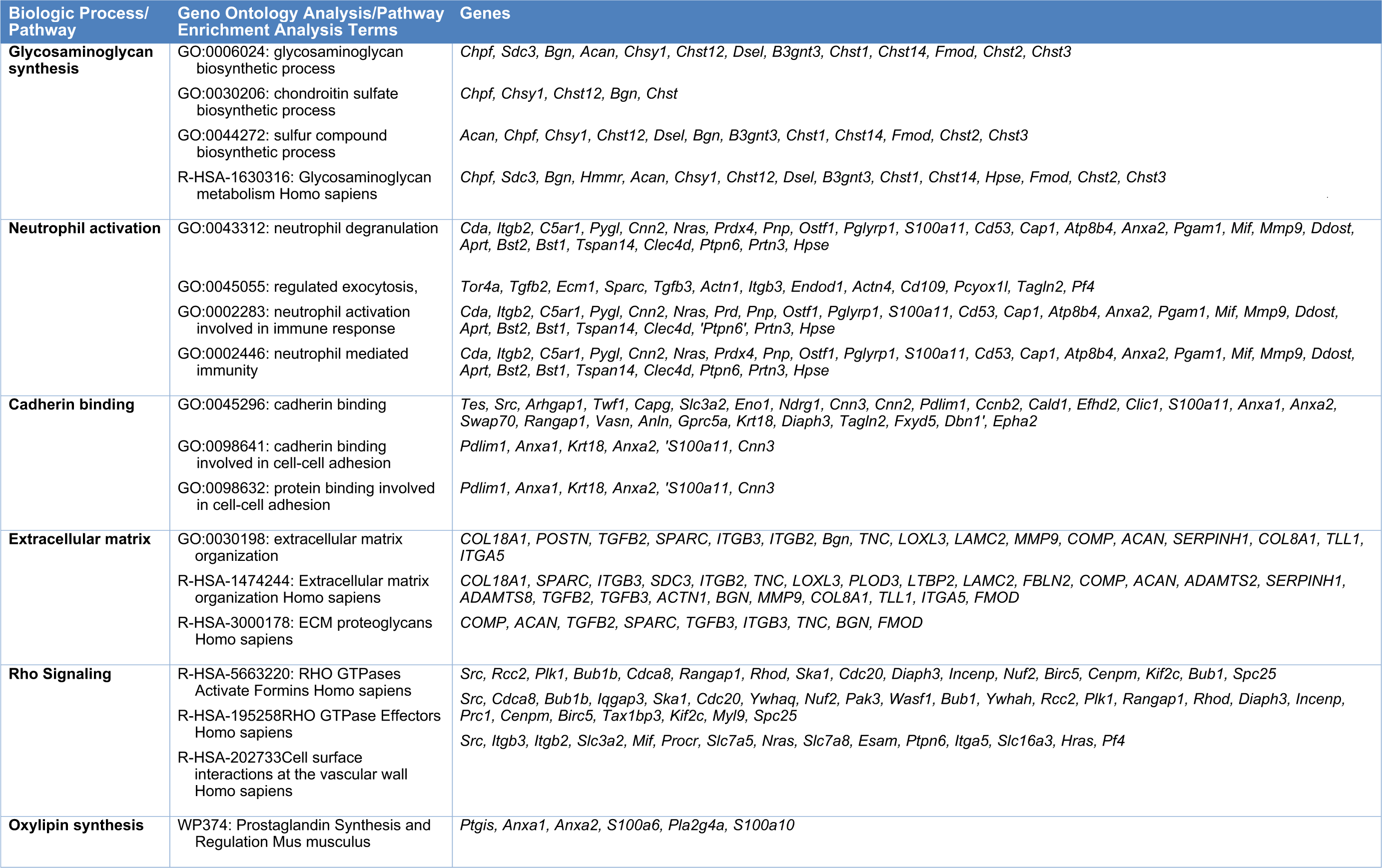
Biological processes augmented in the Ffar4KO infarcted myocardium three days after CAL.

**Table 2:**
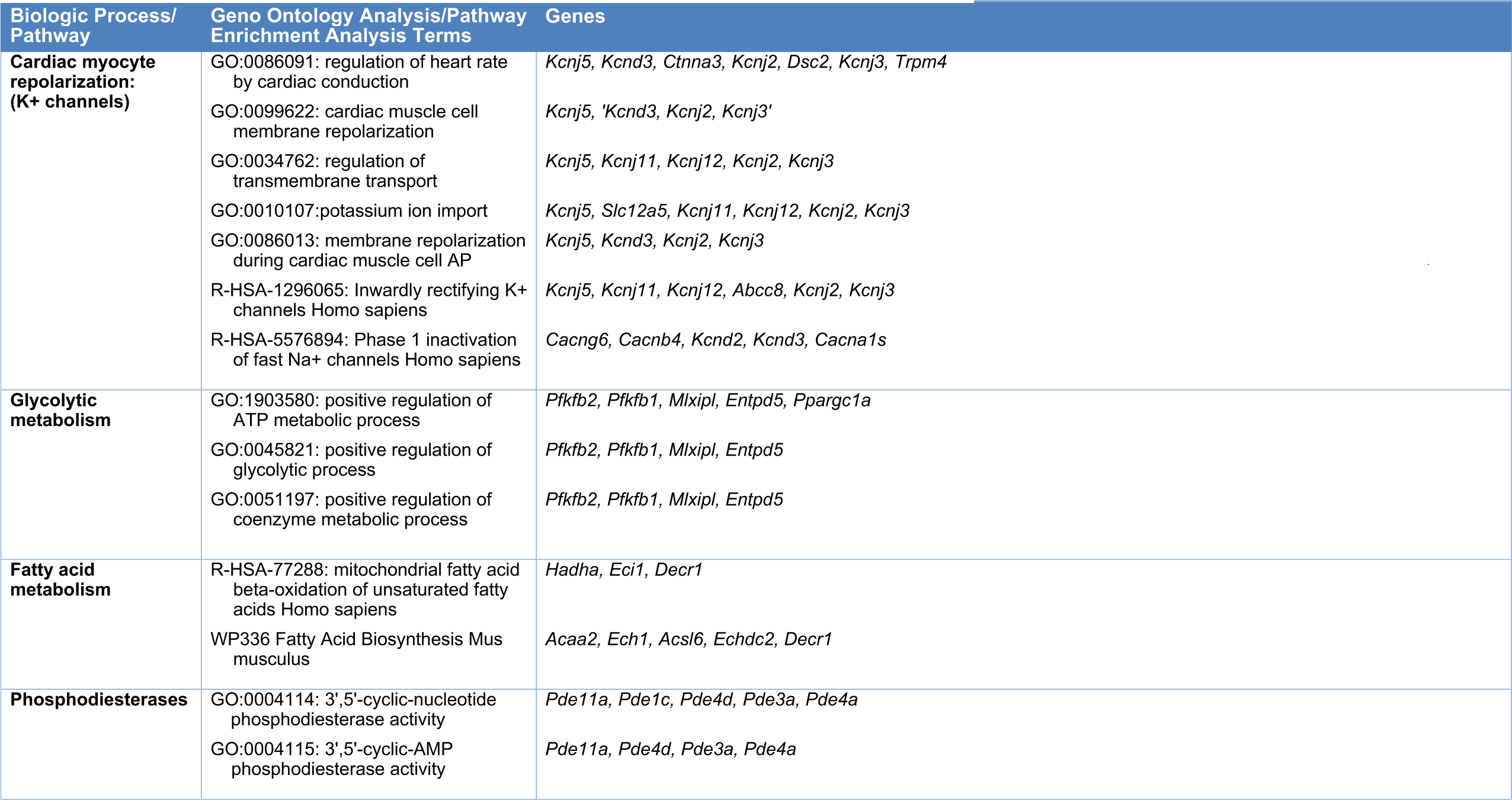
Biological processes impaired in the Ffar4KO infarcted myocardium three days after CAL.

Kyoto Encyclopedia of Genes and Genomes (KEGG) pathway analysis using Pathview ^29^ identified numerous KEGG pathways that were significantly altered in the Ffar4KO infarcted myocardium. Interestingly, AMPK signaling was decreased in the Ffar4KO infarcted myocardium (Figure 4A). Decreased AMPK signaling was defined by a *decrease* in the expression of the following **1.** *Ampk*; **2.** *Adra1A* (α1A-adrenergic receptor), which is notably cardioprotective;^31, 32^ **3.** *Pgc1α* (PPARγ coactivator 1α), which might impair mitochondrial biogenesis.^33^ **4.** *Pfk2* (phosphofructo-2-kinase) and *Fbp* (fructose bisphosphatase-1), which might impair glycolysis; and **5.** *Acc2* (Acetyl-Carboxylase-beta) and *Cpt1* (carnitine palmitoyltransferase-1), which could impair mitochondrial fatty acid β-oxidation. This correlated with the downregulation of glycolytic and fatty acid metabolism identified in the GO and pathway enrichment analyses (Figure 3C, E and Table 2). Conversely, cellular senescence was increased in the Ffar4KO infarcted myocardium (Figure 4B). Increased cellular senescence was indicated by increased expression of *p21* and several cyclins (*CycA*, *CycB*, *CycD*) and cyclin dependent kinases (*Cdk1*, *Cdk4/6*). Senescent cells are also characterized by a senescence-associated secretory phenotype (SASP), and in heart this can include anti-proliferative proteins, pro-inflammatory molecules, matrix metalloproteinases, adhesion molecules and sirtuins.^34, 35^ Analysis of the transcriptome data identified a specific SASP associated with the loss of Ffar4 in the infarcted myocardium (Table 3).

**Figure 4.**
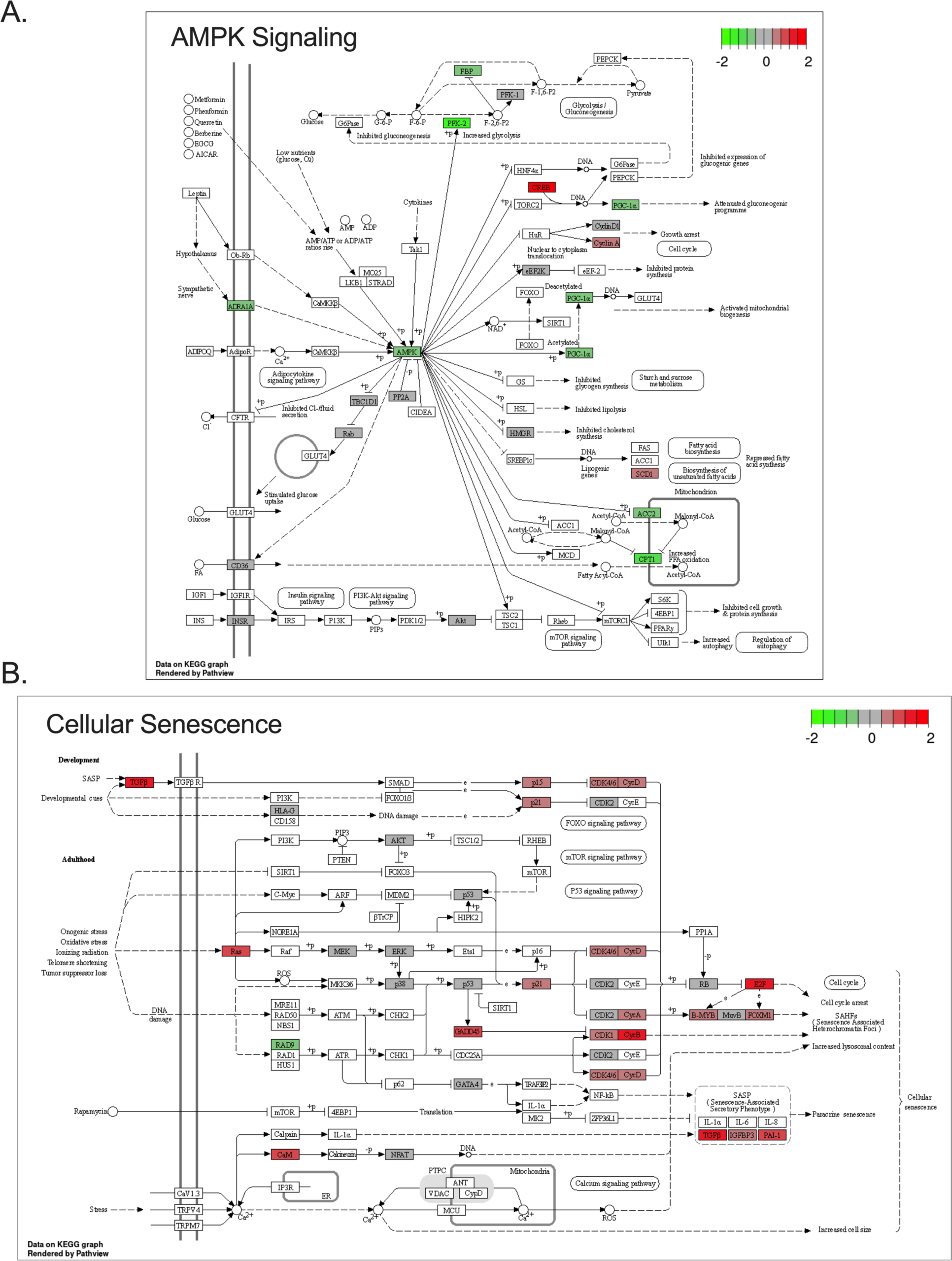
Loss of Ffar4 is associated with decreased AMPK signaling and increased cellular senescence in the infarcted myocardium. Kyoto Encyclopedia of Genes and Genomes (KEGG) pathway analysis of the top 3,000 differentially expressed genes comparing the infarcted myocardium from WT and Ffar4KO hearts was performed using Pathview.^29^ (**A**) The AMPK Signaling pathway was downregulated in the Ffar4KO infarcted myocardium. Downregulated genes included. *Ampk*, *Adra1a*, *Pgc1α*, *Acc2*, *Cpt1*, *Pfk2*, and *Fbp*. (**B**) The Cellular Senescence pathway was upregulated in the Ffar4KO infarcted myocardium. Upregulated genes included: *Tgfb*, *Igfbp3*, *Pai-1* (all 3 part of the senescence associated secretory pathway, SASP), as well as, *p15*, *p21*, *CycA*, *CycB*, *CycD*, *Cdk1*, *Cd64/6*, *Gadd45*, *CaM*, *Ras*, *E2F*, *FoxM1*, and *B-Myb*.

**Table 3.** Specific senescence-associated secretory phenotype associated with Ffar4KO.

| Gene Name | Log2FC | FC | P-Val |
| --- | --- | --- | --- |
| *IL-18* | 0.4726 | 1.3876 | 0.0272 |
| *Cxcl10* | 0.8707 | 1.8285 | 0.0448 |
| *Cxcl3* | 0.8679 | 1.8250 | 0.1632 |
| *Ccl2 (MCP-1)* | 0.9003 | 1.8664 | 0.0193 |
| *MMP9* | 1.0883 | 2.1262 | 0.0011 |
| *Mmp12* | 1.0023 | 2.0031 | 0.0128 |
| *Mmp14* | 0.5102 | 1.4242 | 0.0042 |
| *Sema3F* | 0.4838 | 1.3984 | 0.0047 |
| *Serpine1 (PAI-1)* | 0.9321 | 1.9081 | 0.0069 |
| *Serpine2 (PAI-2)* | 1.3673 | 2.5798 | 0.0635 |
| *Timp1* | 0.6141 | 1.5306 | 0.0772 |
| *Tnfrsf1B* | 0.9094 | 1.8783 | 0.0247 |
| *Tgfb3* | 0.5020 | 1.4162 | 0.0001 |
| *Cpt1* | -0.9327 | 0.5239 | 0.0052 |

In summary, transcriptome analysis of the Ffar4KO infarcted myocardium identified the augmentation of biologic processes (e.g. glycosaminoglycan biosynthesis, neutrophil activation) and signaling pathways (e.g. cellular senescence), along with impairment of biological processes (e.g. glycolytic metabolism) and signaling pathways (AMPK signaling) that would all predict worse cardiovascular outcomes.

### Increased expression of PDE6c attenuates soluble guanylyl cyclase generation of cGMP in Ffar4KO cardiac myocytes

An analysis of the top differentially expressed genes using a volcano plot (Figure 5A) identified 97 genes upregulated (Table 4) and 53 genes (Table 5) downregulated by greater than 2-fold in the infarcted myocardium from male Ffar4KO versus WT hearts. Phosphodiesterase 6C (*Pde6c*) demonstrated the largest fold increase amongst all genes, and it was increased 4.98-fold in the infarcted myocardium from Ffar4KO hearts. Of note, *Pde6c* expression was increased in both cardiac myocytes and non-myocyte fraction isolated from Ffar4KO versus WT hearts (Figure 5B), and Pde6c protein levels were increased in whole heart extracts from Ffar4KO versus WT hearts (Figure 5C-D). Additionally, analysis of a recently published single-cell sequencing analysis of human ischemic cardiomyopathy indicated upregulation of *PDE6C* in ischemic human cardiac myocytes (Figure 5E).^36^ *Pde6c* encodes the α-prime subunit of cone phosphodiesterase, originally named for its expression in the eye, and catabolizes the breakdown of cGMP.^37^ To define the functional consequence of increased *Pde6c* expression in Ffar4KO hearts, primary adult mouse cardiac myocytes were treated with atrial natriuretic peptide (ANP) or vericiguat (Vcgt, soluble guanylyl cyclase agonist) and cGMP levels were measured by ELISA. Interestingly, activation of the natriuretic receptor signaling with ANP had little effect on cGMP levels, whereas activation of soluble guanylyl cyclase with Vcgt significantly increased cGMP levels in WT cardiac myocytes, which was entirely prevented in Ffar4KO cardiac myocytes (Figure 5F). To determine if activation of Ffar4 increases cGMP levels, WT mice were treated for 3 days with the Ffar4 agonist TUG-891, and cGMP was then measured in isolated cardiac myocytes. TUG-891 increased cGMP levels specifically in cardiac myocytes (Figure 5G), suggesting that Ffar4 may regulate cardiac myocyte cGMP to attenuate post-I/R systolic dysfunction.

**Figure 5.**
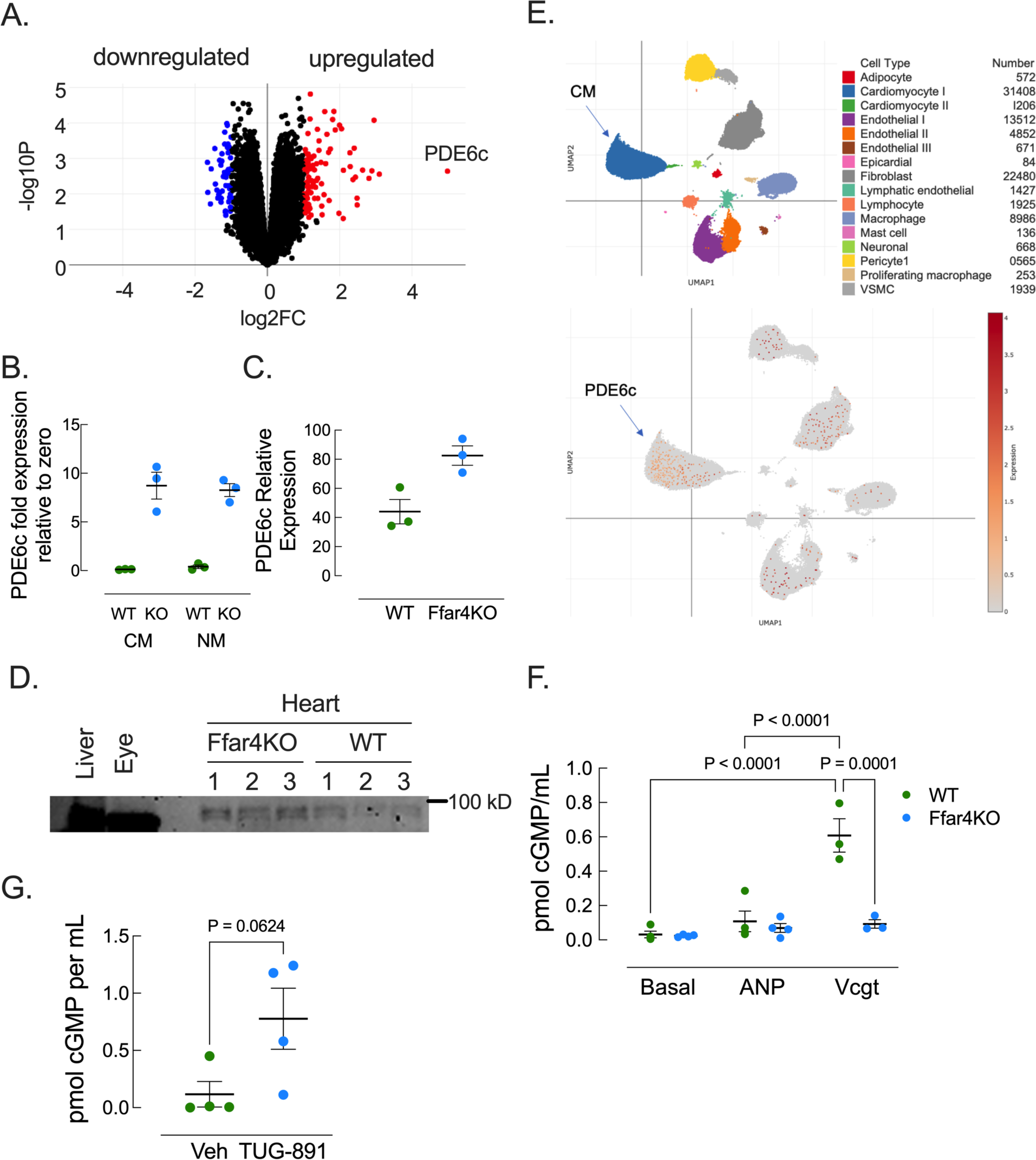
Increased expression of PDE6c attenuates soluble guanylyl cyclase generation of cGMP in Ffar4KO cardiac myocytes. (**A**) Volcano plot showing differentially expressed genes in the Ffar4KO versus WT infarct region, with log2 fold change (cutoff: ±1.0 log2FC) on the x-axis and −log10P on the y-axis (cutoff: P<0.05). (B) PDE6c mRNA expression in WT and Ffar4KO (KO) cardiac myocytes (CM) and non-myocytes (NM). (**C**, **D**) PDE protein expression in WT and Ffar4KO hearts. Western blot of WT and Ffar4KO heart lysates, with liver and eye lysates as positive controls (blot: D; quantitation: C). (**E**) Single-cell sequencing analysis of human ischemic cardiomyopathy identifies PDE6c in cardiac myocytes.^36^ (**F**) Adult cardiac myocytes (CM) were isolated and cultured WT and Ffar4KO hearts. CM were treated with vehicle, atrial natriuretic factor (ANP, 10 μM), or vericiguat (Vcgt, 10 μM) for 30 min and cGMP levels were measured by ELISA. (**G**) WT and Ffar4 mice were treated with vehicle or TUG-891 (35 mg/kg, IP, every 24 h for 3 d), cardiac tissue lysates were prepared and cGMP levels were measured by ELISA. In B, C, F, and G data are expressed as mean±SEM. In B, C, and G, groups were compared by Student’s t-test. In F, data were analyzed by two-way ANOVA with repeated measures with a Tukey post-test to detect differences between groups. P<0.05 was considered significant.

**Table 4:**
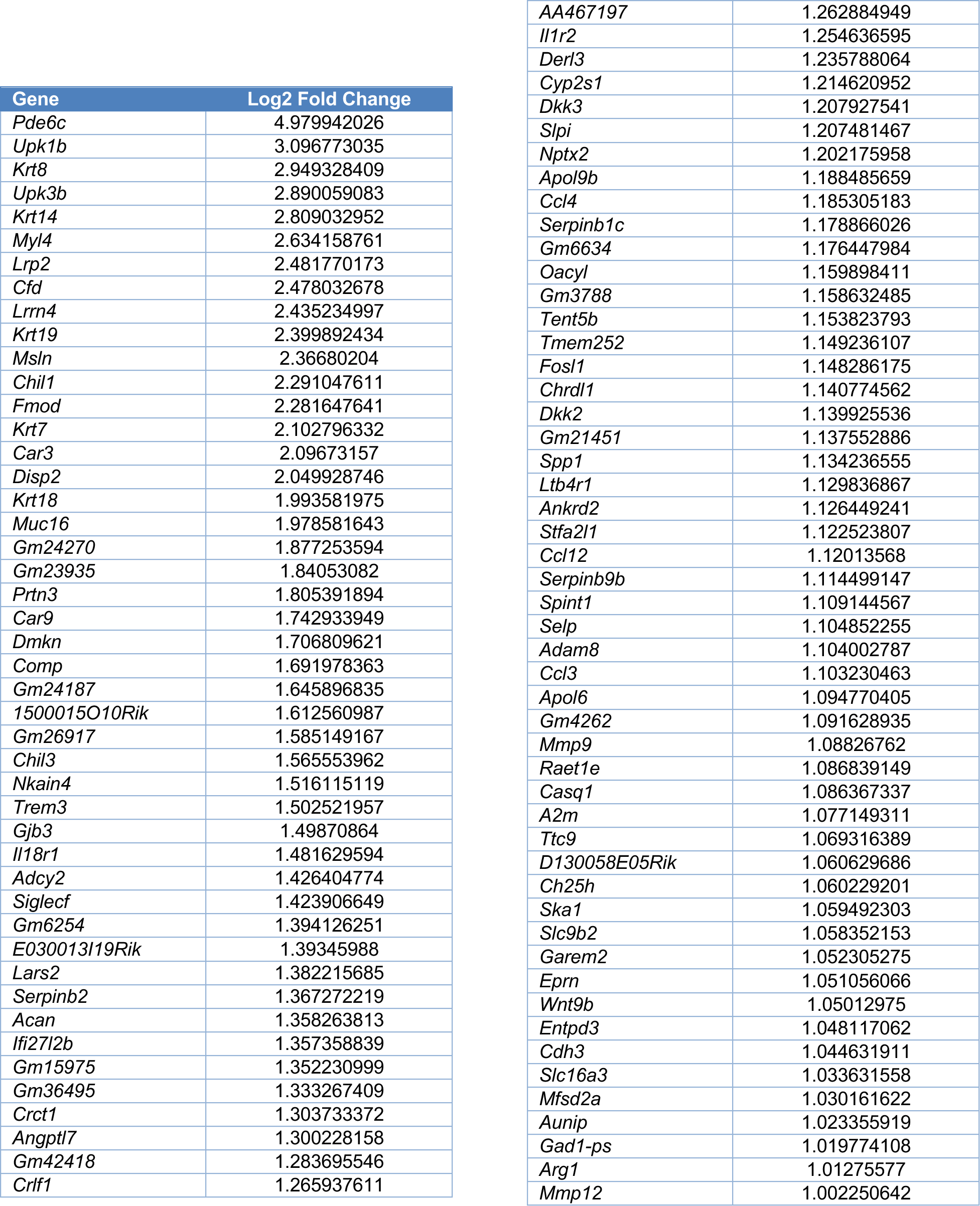
Top upregulated genes in the infarcted myocardium from Ffar4KO Hearts.

**Table 5:** Top downregulated genes in the infarcted myocardium from Ffar4KO Hearts.

| Gene | Log2 Fold Change |
| --- | --- |
| *Lrrc52* | -1.002186141 |
| *Nlrp1c-ps* | -1.00265199 |
| *A330023F24Rik* | -1.006098204 |
| *Erich3* | -1.013900908 |
| *Fbxo47* | -1.015396036 |
| *Gm43481* | -1.018197369 |
| *Gm35339* | -1.01941005 |
| *Gm36936* | -1.021688726 |
| *Gm44949* | -1.022697394 |
| *Uckl1os* | -1.043074113 |
| *Gm15704* | -1.044217466 |
| *Klra9* | -1.069001225 |
| *Gm45137* | -1.069914574 |
| *Gm28151* | -1.070367275 |
| *Pde11a* | -1.076720957 |
| *2900005J15Rik* | -1.077118497 |
| *Gm26621* | -1.084150775 |
| *Gm36670* | -1.086416489 |
| *Gm17276* | -1.088537692 |
| *Gm42208* | -1.095756261 |
| *Cacna1s* | -1.10075014 |
| *Gm43328* | -1.107659004 |
| *Al480526* | -1.107849248 |
| *Nrg2* | -1.108094938 |
| *Myo5c* | -1.121062502 |
| *Xcr1* | -1.135086076 |
| *BC002189* | -1.14409477 |
| *D030034A15Rik* | -1.149080648 |
| *mt-Nd3* | -1.150417966 |
| *9430062P05Rik* | -1.152234397 |
| *Gm33016* | -1.158434719 |
| *Retnla* | -1.162377516 |
| *Gm38077* | -1.17821153 |
| *Gm20342* | -1.179567228 |
| *A930004J17Rik* | -1.188288804 |
| *Gm17745* | -1.21705493 |
| *Lgals4* | -1.247038628 |
| *Gm43185* | -1.256758777 |
| *Gm43858* | -1.259063448 |
| *Clec4b1* | -1.276921165 |
| *Cfap61* | -1.279831719 |
| *Gm17428* | -1.284609204 |
| *Gm40652* | -1.320149799 |
| *Gm13479* | -1.328001124 |
| *Gm37893* | -1.341356793 |
| *Gm14964* | -1.372736767 |
| *Gm40604* | -1.396937745 |
| *Gm49405* | -1.439335084 |
| *Gm17024* | -1.455807768 |
| *Aldob* | -1.555341898 |
| *A930017K11Rik* | -1.576491388 |
| *Gdap10* | -1.647840923 |
| *4632404M16Rik* | -1.65945464 |

### Cardiac myocyte-specific overexpression of Ffar4 attenuates left ventricular systolic dysfunction post-I/R in male mice

The worsened systolic function induced by I/R in Ffar4KO mice without an effect on left ventricular remodeling suggests that the loss of Ffar4 specifically in cardiac myocytes might explain the worsened left ventricular systolic function observed in the Ffar4KO mice. Therefore, to test the hypothesis that activation of Ffar4 in cardiac myocytes would attenuate I/R induced left ventricular systolic dysfunction, we quantified left ventricular remodeling in male wild-type (WT) mice and mice with cardiac myocyte-specific overexpression of Ffar4 (CM-Ffar4-Tg) subjected to 120 minutes of ischemia by transient coronary artery ligation followed by reperfusion for four weeks. The duration of ischemia was increased to 120 mice with the goal of inducing more cardiac injury, and thus decreasing the rate of systolic functional recovery observed with 60 minutes of ischemia in WT mice. Male mice were tested because no sex differences were identified in the original knockout experiment (Figures 1-2). Ffar4 transgene expression in the CAG-CAT-Ffar4 mice was induced by AAV9-cTnt-Cre virus four weeks prior to I/R surgery, with AAV9-cTnt-GFP used as a control (referred to as ‘WT mice’). Ffar4 expression was quantified in WT (AAV9-cTnt-GFP) and CM-Ffar4-Tg (AAV9-cTnt-Cre) hearts four weeks following virus injection in a subset of mice. In isolated cardiac myocytes and non-myocytes, Ffar4 expression was specifically increased three-fold in cardiac myocytes isolated from CM-Ffar4-Tg hearts, but not in the non-myocyte fraction (Figure 6A). Three days after surgery, I/R induced a significantly greater decrease in left ventricular systolic function in male WT versus CM-Ffar4-Tg mice (EF, Figure 6B-D). From that point forward, systolic function recovered to almost baseline values in male CM-Ffar4-Tg mice by four weeks, but remained suppressed in male WT mice (Figure 6B). Left ventricular weight was not different in WT versus the CM-Ffar4-Tg hearts after four weeks (Figure 6E). These results indicate that Ffar4 in cardiac myocytes specifically attenuates left ventricular systolic dysfunction post-ischemia in male mice.

**Figure 6.**
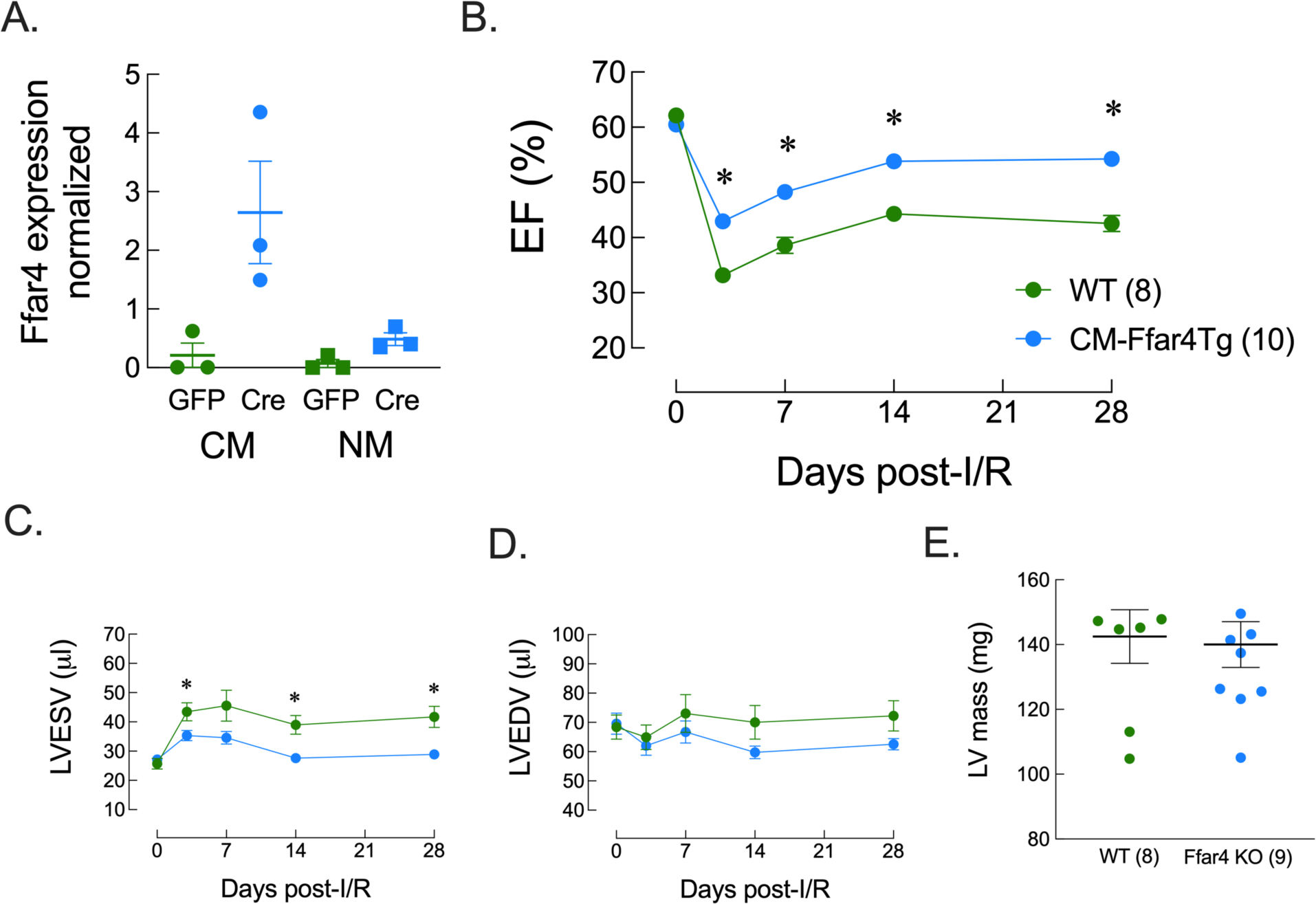
Cardiac myocyte-specific overexpression of Ffar4 attenuates left ventricular systolic dysfunction post-I/R in male mice. (**A**) To quantify the level of Ffar4 expression in transgenic mice, mice with the CAG-lox-stop-lox-CAT-Ffar4 construct were injected with AAV9-cTnt-<u>GFP</u> control virus (WT) or AAV9-cTnt-<u>Cre</u> virus (Ffar4Tg) at 8 weeks of age. After 4 weeks, cardiac myocyte (CM) and non-myocytes (NM), total RNA was isolated, and Ffar4 expression was quantified by qRT-PCR. In a separate cohort of mice, AAV9-cTnt-GFP control virus (WT) or AAV9-cTnt-Cre virus (Ffar4Tg) was delivered to male CAG-lox-stop-lox-CAT-Ffar4 mice at 8 weeks of age. Four weeks after viral delivery, mice were subjected to 120 min of ischemia followed by reperfusion for 4 weeks. Systolic function was measured as in Figure 1. (**B**) Ejection Fraction, EF; (**C**) Left ventricular end diastolic volume, LVEDV; (**D**) Left ventricular end systolic volume, LVESV; and (**E**) Left ventricular mass. In A-E, data are expressed as mean±SEM. In B-D, data were analyzed by two-way ANOVA with repeated measures with a Tukey post-test to detect differences between groups. In E, data were analyzed by Student’s t=test. P<0.05 was considered significant.

## Discussion

Here, we have defined a novel cardioprotective role of Ffar4 to prevent ischemic cardiomyopathy. We found that loss of this receptor for long chain fatty acids, including cardioprotective μ3-PUFAs, impaired the recovery of left ventricular systolic function in male and female mice post-I/R despite no initial difference in infarct size (Figures 1-2), suggesting that Ffar4 is required for an adaptive response to I/R. Furthermore, we demonstrated that overexpression of Ffar4 specifically in cardiac myocytes is sufficient to attenuate left ventricular systolic dysfunction post-I/R, identifying for the first time a cardiac myocyte-specific cardioprotective role for Ffar4.

Transcriptomic analysis comparing the infarcted myocardium from WT and Ffar4KO mice three days post-I/R identified several potential mechanistic explanations for the cardioprotective function of Ffar4. In the Ffar4KO infarcted myocardium, pathway analysis indicated augmentation of glycosaminoglycan synthesis (chondroitin sulfate), neutrophil activation, cadherin binding, extracellular matrix, rho signaling, and oxylipin synthesis (Figure 3, Table 1). Conversely, pathway analysis revealed downregulation of glycolytic and fatty acid metabolism, cardiac myocyte repolarization (K+ channel function), and phosphodiesterase, activity in the Ffar4KO infarcted myocardium (Figure 3, Table 2). Further, KEGG pathway analysis indicated reduced AMPK signaling and increased cellular senescence in the Ffar4KO infarcted myocardium (Figure 4). Each of these pathways individually might explain the worsened outcomes post-I/R in the Ffar4KO mice, but the potential exists that some of these pathways are interconnected. In simplified terms, the sequalae of post-I/R remodeling in cardiac myocytes involves mitochondrial damage and increased oxidative stress, leading to cardiac myocyte death (necrosis, apoptosis, pyroptosis), release of DAMPs, activation of PRRs and immune cell infiltration.^3–5^ I/R-induced mitochondrial damage and release of DAMPs in cardiac myocytes might also provoke increased NLRP3 inflammasome activity leading to pyroptosis.^38^ Several studies in other cell types suggest that Ffar4 might affect these pathways in cardiac myocytes. For instance, Ffar4 increases mitochondrial metabolism in brown fat,^39^ decreases mitochondrial ROS,^16, 17^ and induces mitophagy to protect against oxidative stress.^18^ Further, Ffar4 activates AMPK to regulate cholesterol efflux in macrophages,^40^ as well as prevent steatosis^41^ and alcoholic liver disease.^42^ Ffar4 induces an M2-like, pro-resolving phenotype in macrophages.^8^ Additionally, Ffar4 attenuates activation of the NLRP3 inflammasome in Kupffer cells^19^ and in macrophages.^20^ Ffar4 activation of an APMK/Sirt3 signaling pathway attenuates senescence in kidney epithelial cells.^15^ Based on this, we hypothesize that Ffar4 might protect cardiac myocytes, at least in part, by activating AMPK signaling to reduce oxidative stress, thereby preserving mitochondrial function, but proving this will require more detailed mechanistic studies in cardiac myocytes.

Interestingly, the most differentially expressed gene in the Ffar4KO mice was PDE6c (Figure 5A-D), which is uncharacterized in heart, but is significantly increased in cardiac myocytes in human ischemic cardiomyopathy (Figure 5E).^36^ Functionally, we demonstrated that the soluble guanylyl cyclase stimulator vericiguat significantly increased cGMP in WT cardiac myocytes, but showed no effect in Ffar4KO cardiac myocytes, consistent with an increase in PDE6c expression. Cardiac myocyte cGMP levels are spatially and temporally regulated through NO-dependent activation of soluble guanylyl cyclases (sGC) or natriuretic peptide receptor activation of particulate GC (GC-A, GC-B), and breakdown by phosphodiesterases.^43^ Clinically, targeting cGMP signaling to improve HF outcomes has met with mixed success.^44^ In the VICTORIA trial, vericiguat reduced cardiovascular death and hospitalization for heart failure by 10% in patients with heart failure with reduced ejection fraction.^45^ Conversely PDE5 inhibitors, such as sildenafil, have not improved mortality or decreased heart failure hospitalizations in clinical trials.^46^ Increased PDE6c activity might partly explain these failures. The IC50 of sildenafil for PDE5 is 3.5nM but for PDE6 it is 33nM (i.e. only 10% as effective on PDE6).^47^ Therefore, it is possible that lack of inhibition of PDE6 activity by sildenafil resulted in suboptimal inhibition of cGMP hydrolysis in heart failure trial participants. Finally, we note that the Ffar4 agonist TUG-891 increased cardiac cGMP levels, which might suggest that activation of cGMP signaling is part of the benefit of Ffar4 post-I/R that was observed in the CM-Ffar-Tg (Figure 6).

Therapeutically, targeting Ffar4 with μ3-PUFAs might have clinical utility. Many clinical trials have shown that μ3-polyunsatured fatty acids reduce mortality in CHD,^48–52^ while one large clinical trial^53^ and several smaller trials^54–56^ also indicate that μ3-PUFAs improve outcomes in HF. Yet, some clinical trials with μ3-PUFAs in CHD have been null,^57–59^ which might have more to do with trial design than μ3-efficacy.^60^ Interestingly, trials with high-dose eicosapentaenoic acid (EPA), JELIS (EPA, 2.8 g/d) and REDUCE-IT (Icosapent Ethyl, 4g/d), reported significant reductions major adverse cardiovascular events.^61, 62^ Furthermore, high-dose μ3-PUFAs attenuated post-infarct remodeling after 6 months in participants in the OMEGA-REMODEL trial indicated by decreased LV end-systolic volume index, non-infarct fibrosis, infarct size, and serum markers of inflammation, which all positively correlated with μ3-index and EPA enrichment.^63^ However, the negative results of STRENGTH with high-dose μ3-PUFAs (4g Epanova, EPA and docosahexaenoic acid (DHA) μ3-carboxylic acids) might temper the positive outcomes with high-dose EPA.^64^ In total, existing clinical data suggests that the beneficial effects of μ3-PUFAs, particularly EPA, in CHD and HF may be concentration-dependent and thus receptor-mediated.

In summary, we have shown that Ffar4 in cardiac myocytes attenuates ischemic cardiomyopathy, and suggested several potential mechanisms to explain this benefit, including reduction in oxidative stress, preservation of mitochondrial function, and modulation of cGMP signaling. Finally, our findings have conceivable translational benefits given the known efficacy of μ3-PUFAs to improve outcomes post-MI in humans consistent with those observed in the OMEGA-REMODEL trial.

## Acknowledgements

The authors acknowledge the University (of Minnesota) Imaging Centers (UIC) for support on echocardiography.

## Author Contributions

Conceptualization: MJZ, GCS, TDO

Methodology: MJZ, SK, DJG, SP, CLH, SCW, GCS, TDO

Formal analysis: MJZ, SK, SP, TDO

Investigation: MJZ, SK, DJG, SP, CLH, SCW, GCS, TDO

Writing-Original Draft: MJZ, TDO

Writing-Review & Editing: MJZ, SK, DJG, SP, CLH, SCW, GCS, TDO

Visualization: MJZ, TDO

Supervision: GCS, TDO Funding Acquisition: GCS, TDO

